# Production and control of a Brazilian tailor-made meningococcal B vaccine

**DOI:** 10.1101/2021.08.13.456295

**Authors:** E Jessouroun, AP Santos, ES Figueira, MB Correa, ML Leal

## Abstract

Meningococcal disease has been a public health problem in Brazil since the serogroups A and C epidemics occurred in the 1970s. The Oswaldo Cruz Foundation in Brazil has been working to develop a serogroup B meningococcal vaccine composed of detergent-treated outer membrane vesicles (OMV) and detoxified endotoxin (dLOS) from *Neisseria meningitidis* serogroup B prevalent strains. Experimental vaccine were produced in a pilot-scale under Good Manufacturing Practice condition (GMP). Physicochemical and biological controls were established based on what has been reported previously for OMV vaccines. The developed vaccine contained the main class 1, 2, 3, and 5 proteins, some minor iron-regulated proteins, and 5–10% residual lipopolysaccharide related to total protein content. The dLOS was added, as a vaccine component, in half of the total protein amount. The pyrogenicity of the final products, based on the residual LOS in OMVs, varied from 1.01 to 2.24 ng LOS/kg rabbit. Three experimental vaccines, were highly immunogenic in mice with better performance using higher antigen concentrations.

## INTRODUCTION

Meningococcal disease remains a worldwide health problem and has been a public health problem in Brazil since the groups A and C epidemics of the 1970s. Serogroup B caused 80% of the meningococcal disease cases throughout the 1980s with a significant number of cases in children < 2-years-of-age [1,2].

The, B:4,7:P1.19,15, L3,7,8 phenotype was responsible for the 1988 São Paulo epidemic, continued to represent the predominant B phenotype from 2000 to 2004, and seems to be the main strain today [3,4].

The other prevailing phenotypes identified were B:4,7:P1.7, L3,7 and B:15:P1.7,16, which have high frequencies in the southern states of Brazil. These three prevalent serosubtypes belong to the same ET-5 electrophoretic type [5].

The emergence of W135, which is associated with the ST-11 complex, was reported in São Paulo, Rio de Janeiro, and Rio Grande do Sul. Serogroup C was responsible for 75% of the cases identified in 2010, 17% by serogroup B, 6% by serogroup W135, and 2% for serogroup Y [1].

The Brazilian Ministry of Health decided the use a Cuban vaccine (VAMENGOC-BC^®^) in the southern region of Brazil in the 1990s when they were trying to control the meningococcal B epidemic. Case control studies to estimate vaccine efficacy in Brazil were performed from 1989 to 1990. The results indicate that the vaccine generally did not provide protection against heterologous strains and only induced modest protection in young children [6,7,8].

Based on the results of the use of the Cuban vaccine in Brazil, three Brazilian research institutions (Bio-Manguinhos/Fiocruz, Institutos Butantan, and Adolfo Lutz) have collaborated to develop a serogroup B meningococcal vaccine composed of detergent-treated OMVs and detoxified lipooligosaccharide (dLOS) from Brazilian prevalent strains. The Brazilian approach has focused on developing a tailor-made vaccine that have been developed and successfully proven efficacious against serogroup B meningococcal disease in Cuba, Norway, and New Zealand. The inclusion of dLOS as a vaccine component, was based on the fact that other subcapsular antigens, in addition to OMV, may benefit cross-immunitity within different serotypes of meningo B strains [9,10].

Preliminary pre-clinical data showed animal protection induction of experimental vaccines from an intraperitoneal challenge model in mice, and protective bactericidal antibody against vaccine strains [11].

The study presented here describes a vaccine standardized production and control under Good Manufacturing Practice (GMP) conditions in a pilot-scale process. The results indicate production and controls compatible with OMV vaccines and potential use of the developed product for clinical studies for safety and immunogenicity assessment [12,13].

## MATERIAL AND METHODS

### Vaccine preparation

#### Vaccine strains

The N44/89 (B:4,7:P1.19,15, L3,7,9) and N603/95 (B:4,7:P1.7.1, L3,7) strains were provided by the Adolfo Lutz Institute, São Paulo, Brazil.

#### Biomass harvest

The production process included cultivation of the vaccine strains in a 150 L bioreactor (B Braun Model Biostat UD 100) using Catlin MC.6 with 20 μM Fe^+3^ and 42 μM ethylene diamine-di-(*O*-hydroxy-phenylacetic acid) (EDDHA) to produce the OMVs. Lipooligosaccharide (LOS) was produced in the same way described for OMVs without EDDHA. Bacterial growth was inactivated with heat (56°C/30 min), and the biomass was separated by continuous flow centrifugation (LAPX 4045GP-31G; Alfa Laval) [14].

#### OMV antigens

The LOS-depleted OMVs were obtained from the culture supernatant, and concentrated by tangential ultrafiltration in a Centrasette stainless steel cassette holder using a polyethersulfone Centrasette cassette (0.9-m^2^) and a molecular weight cutoff of 100 kDa (Pall Corp., Port Washington, NY, USA). The concentrate was treated with 2% (w/v) sodium desoxycholate (DOC) and LOS-depleted OMVs were harvested by ultracentrifugation (100,000*g* for 2 h), in water for injection containing 3% sterile-filtered sucrose using a 0.22-μm Millipak^®^-200 polyvinylidene fluoride membrane (PVDF; Millipore, Billerica, MA, USA) [11].

#### Detoxified LOS antigen

LOS from N44/89 strain, was obtained from the culture biomass after hexadecyl trimethyl ammonium bromide (Cetavlon-Fluka Analytical code 52370) treatment using the hot phenol/water method (Gotschlich,1969; Westphal,1965). The extracted LOS was further purified by gel filtration chromatography using 20 mM Tris-HCl, pH 8.5, containing 0.5% DOC and 5 mM EDTA as the mobile phase on a single Sephacryl HR S-300 column (ID: 5.0 μm L: 100.0 cm). The LOS sample solution applied to the column was limited to 4% of bed volume and a LOS mass not greater than 600 mg [15].The detoxified LOS (dLOS) was obtained by treatment of raw material with 0.20 M NaOH in a water bath at 60°C for 150 min [16,17].

#### Vaccine formulation

Vaccine 1: [25 μg OMV protein (12.5 μg from N44/89 +12.5 μg from N603/95 + 12.5 μg dLOS from N44/89) + Al(OH)_3_/2 mg/mL].Vaccine 2: [50 μg OMV protein (25 μg from N44/89 + 25 μg from N603/95 + 25 μg of dLOS from N44/89 + Al(OH)_3_/2 mg/mL]. Vaccine 3: [100 μg OMV protein (50 μg from N44/89 + 50 μg from N603/95 + 50 μg of dLOS from N44/89 + Al(OH)_3_/2 mg/mL]. The final quality control tests were performed in the Quality Control Department of Bio-Manguinhos (Rio de Janeiro, Brazil). Vamengoc BC^®^ and DTP vaccine as the reference in pyrogen test.

### Vaccine response studies

#### Study Groups

Swiss Webster male and female mice (weight, 12–17 g) were obtained from Oswaldo Cruz Foundation breeding unit (Rio de Janeiro, Brazil) and caged with free access to food and water in a room maintained at 22–24°C with a 12 h:12 h light/dark cycle in the Bio-Manguinhos Experimental Animal Facility. The animal procedures were conducted according to the Oswaldo Cruz Foundation Ethical Committee for animal using (CEUA/FIOCRUZ number LW65/14). Groups of 15 mice were immunized intramuscularly with 0.2 mL of the final experimental product (diluted 1:10 in phosphate buffered solution (PBS). A three-dose immunization schedule of 14 days intervals was used. Groups of five mice were bled (retro-orbital via) before each dose and 15 days after the third immunization.

#### OMV-enzyme-linked immunosorbent assay (ELISA)

To detect anti-OMV total IgG, 96-well plates (ref. no. 3590; Corning-Costar, Corning, NY, USA) were coated with 100 μL/well of OMV (4 μg/mL) from N44/89 or N603/95 strains. Sera samples were diluted in two-fold serial dilutions in TBS with 0.05%Tween 20 (Merck, Schuchardt, Germany) and 5% FCS (Sigma-Aldrich, St. Louis, MO, USA) and incubated overnight at 4°C. The plates were incubated with anti-mouse IgG conjugated with alkaline phosphatase (whole molecule) (Sigma-Aldrich A-3688) diluted 1:2000. All sera were titrated in duplicate, and the titers were determined at an absorbance of 405 nm using a VERSA max tunable microplate reader (Molecular Devices, Sunnyvale, CA, USA). As the antibody standard, a positive post vaccination serum was used in all experiments. The observed optical density was transformed to arbitrary units per milliliter by a sigmoidal standard curve (logit-log transformation) calculated from the values (1000 EU/mL) of the reference serum [18].

#### LOS ELISA assay

Microtiter plates (Immulux ref 1000; Dynex Technologies, Chantilly, VA, USA) were coated overnight at room temperature with 100 μL/well of 10 μg/mL LOS-DOC micelles dissolved in PBS with 0.1% DOC. TBS with 5% bovine serum albumin (BSA) (Sigma-Aldrich ref A9418) was used as the blocking buffer for 2 h at 37°C. Sera samples were titrated in duplicate in TBS with 5% BSA and incubated for 3 h at 37°C. Conjugated antibodies (Sigma-Aldrich A-3688) were diluted 1:1000 and incubated for 2 h at 37°C. The reaction was developed for 20 min with a phosphatase substrate (Sigma-Aldrich S0942). All sera were titrated in duplicate and titers were determined with absorbance at 405 nm in VERSAmax tunable microplate reader – Molecular Devices. The total IgG antibody responses (EU/mL) against OMVs and LOS, were presented as geometric mean concentrations (GMC) of five serum samples (pool of five mice) by using a 4-parameter logistic curve-fitting analysis with SoftMaxPro software and documentation - Molecular Devices, against the standard curve generated by in-house serum with 1000 EU/mL of arbitrary units. All p-values were calculated with Kruskall Wallis test. Differences were considered as statistically significant at *p*< 0.05 [18,19].

#### Serum bactericidal assay (SBA)

The bacterial strains, was streaked on Columbia blood agar (Merck 1.10455.0500), plated with 5% horse blood (CBA), and incubated overnight at 37°C in 5% CO_2_. The next morning, 10–20 colonies, were subculture on another CBA plate and incubated for 4 h at 37°C in 5% CO_2_. After 4 h, the bacteria were suspended in a bactericidal buffer (Hanks balanced salt solution) containing 0.5% BSA (Sigma, Poole, UK) and 0.5 U/mL heparin (CP Pharmaceuticals, Wrexham, UK). The bacteria concentration was adjusted to 2 × 10^5^ colony forming units (cfu)/mL. Equal volumes (10 μL) of the bacterial suspension and human complement were added to 20 μL heat-inactivated test serum serially diluted two-fold in bactericidal buffer and added to 96-well U-bottom microtiter plates (Greiner, Frickenhausen, Germany). Human plasma from a suitable donor (Research Ethics Committee, Hemorio Hospital, Rio de Janeiro, Brazil No. 118/07) was used as the complement source. The plasma was used in the assay after a 60 min incubation at 25% of the final concentration. The number of bacteria was determined at time zero, by allowing 10 μL of the reaction mixture to flow 8–10 cm in lanes down a CBA plate (the tilt method). After a 60-min incubation of the reaction mixture at 37°C, 10 μL was removed from each well and plated on CBA. The colonies were counted after a 37°C overnight incubation in 5% CO_2_. SBA titers were expressed as the reciprocal of the final serum dilution step that produces a 50% mortality after 60 min compared to the number of bacteria at time zero. Three in-house control sera (low, medium, and high titers) were used as controls for the assays in each target strain. The test was performed using vaccine strains as the target [20,21].

#### TEST FOR TOXICITY

The experimental vaccines were tested by intraperitoneal injection of one human dose, but not more than 1 mL of experimental vaccine into 5 mice (weight, 17–22 g) diluted in diluent. Two guinea pigs (weight, 250–350 g) were also injected (i.p.) with the equivalent of 10 human doses in a volume < 5 mL. The vaccines tested were considered innocuous if the animals survived at least 7 days without showing significant toxic signs. Cuban (VAMENGOC-BC^®^) and DTP (Butantan Institute, São Paulo, Brazil) were used as the reference of licensed vaccines for human use, in Brazil [22].

#### Rabbit pyrogen test

Briefly, the tested vaccines in various human-dose dilutions were given intravenously to three rabbits (weight, 2.5–3.0 kg) at a rate of 1 mL/kg of rabbit body weight. Cuban (VAMENGOC-BC^®^) and DTP (Butantan Institute, São Paulo, Brazil) were used as a reference of licensed vaccines for human use, in Brazil. Rectal temperature was measured with an indwelling rectal thermometer and recorded 5 h after injecting the sample. The preparations were considered approved if the summed temperature variations of the three rabbits did not exceed 1.15°C [22].

#### Sterility Test

According to the European Pharmacopoeia, eight vials of each vaccine containing five doses each in Al(OH)_3_ vaccine diluent, pooled, and filtered through a 0.22-μm PVDF membrane to fluidize the thioglycolate (TGC) and soy-bean casein media (TSB). The TGC and TSB were incubated at 30–35°C for 24–48 h and 20–25°C for 10–15 days, respectively [23].

### PhysicoChemicalControls

#### Protein Concentration

Briefly, the OMVs were mixed in the copper reagent (4% CuSO_4_·5H_2_O) with sodium dodecyl sulfate (SDS) (1% w/v) for 10 min. Sample absorption was measured after adding Folin Ciocalteu’s phenol reagent and incubating for 45-min at room temperature. Protein content was determined in Beckman DU530 spectrophotometer at 660 nm against a BSA standard curve (Pierce, Rockford, IL, USA) [24].

#### Sodium Dodecyl sulfate-polyacrylamide gel electrophoresis (SDS-PAGE) Pattern

The protein profile of the OMV preparations, before vaccine formulation,was analyzed by 12% PAGE in the presence of 2% (w/v) SDS. The proteins were stained with 0.1% (w/v) Coomassie Brilliant Blue, and analyzed by densitometry. The stained class protein bands were determined by scanning the gel, the relative amount of each protein was calculated using Quantity One software (densitometer GS-800; Bio-Rad Laboratories, Hercules, CA, USA), and was presented as a percentage of total protein [25].

#### LOS and dLOS electrophoresis Pattern

The LOS samples (LOS and dLOS) were treated for 5 min at 100°C in 0.05 M Tris-HCl buffer (pH 6.8), 2% (w/v) SDS, 10% (w/v) sucrose, and 0.01% bromophenol blue and fractionated on an SDS-polyacrylamide gel (8 × 7.3 cm by 0.75 mm) containing 4% and 16.5% acrylamide in the stacking and separating gels, respectively. The electrophoresis was performed at 15 mA in the stacking gel and 20 mA in the separating gel until the tracking dye had traveled about 10 cm. The SDS-PAGE fractionated LOS preparation was stained using the conventional silver staining method of Tsai and Frasch, 1982. A protein standard (16,949–2,512 kDa) was used to estimate the molecular weights of LOS and dLOS. The gel was stained with 0.1% (w/v) Coomassie Brilliant Blue [26]. HP Scanjet G4050 Scanner was used, to scan the gel, with the following parameters: Output Type: Gray Scale; File Type: Unzipped Tif (* .tif) image; Output Resolution: 1200 dpi (dot per inch). The gel image file was analyzed using the Image Master 1D Prime V3.01 software (Pharmacia Biotech, San Francisco, CA, USA)

#### Structural Characterization of dLOS

Detoxified LOS, which is *O*-deacylated (dLOS), was hydrolyzed in acetate buffer at pH 4.5. It was purified by gel-filtration chromatography using a Bio-Gel P-4 column. The dLOS electrospray ionization/mass spectroscopy analysis was performed with a Thermo Finnigan LCQ Deca XP ion trap mass spectrometer (ThermoScientific, Waltham, MA, USA). Capillary temperature was 200°C, capillary voltage was 45 KV, and mass range m/z was 250–2000 [27].

#### Residual LOS in the OMV Preparations

LOS was quantified by determining 2-keto-3-deoxyoctonic acid (KDO) content using a mass conversion factor value of 20. OMVs from each strain, were diluted in 1 M Tris-HCl, 0.5% SDS, and 0.05 M CaCl_2_ buffer, at pH 7.5 following incubation for 5 min at 100°C. The protein in this suspension was digested with proteinase K for 4 h at 56°C. KDO content was measured using the thiobarbituric acid procedure. Briefly, a standard aqueous (0.1 mg/mL) solution of KDO (Sigma ref K2755) was prepared and stored at–20°C. This standard solution was diluted to prepare five different concentrations. The digested samples were hydrolyzed in trifluoracetic acid and treated in sequence with periodic acid, sodium arsenite, and thiobarbituric acid and heated in a boiling water bath for 10 min. KDO content was determined by measuring optical density in a Beckman DU530 spectrophotometer (Beckman, Brea, CA, USA) at 548 nm against a KDO standard curve [28].

##### pH

pH was determined in each experimental vaccine in its diluent according to the United States Pharmacopeia using standard buffer solutions from the Physicochemical Quality Control of Bio-Manguinhos [23].

##### Sucrose content

The quantity of sucrose in the experimental vaccines diluted in water was analyzed by high-performance liquid chromatography (HPLC; Waters, Milford, MA, USA) using a Shodex Sugar SC1011 column (300 × 8 mm) at 75°C and degasified Milli-Q water as the mobile phase at a flow rate of 0.8 mL/min. The sucrose amount was evaluated with a refractive index detector (Waters IR 410 Channel 2) [29].

##### Aluminum hydroxide adsorption

The experimental vaccines were centrifuged (4,000 rpm/20 min/4°C), and protein content was measured in the supernatant. The test was verified by > 80% adsorption of the total vaccine protein concentration [22].

##### Electron microscopy

Droplets of LOS-depleted OMVs from the N44/89 strain (1 mg protein/mL) in distilled water were applied to glow-discharged, carbon-filmed grids for 1 min and negatively stained with 0.5% (w/v) phosphotungstic acid, pH 7.0 for 1 min. The preparations were examined under an electron microscope operated at 100 keV (JEM 1010; Jeol, Tokyo, Japan) [30].

##### Sialic acid content

Residual sialic acid was quantified by HPLC using a pulsed amperometric detector. Before HPLC, the OMV DOC treated, was hydrolyzed with 0.2 M HCl at 100°C for 60 min. HPLC analyses were performed on a Dionex I.C.S.–3000i, with a LC50 gradient pump and an ED50 electrochemical detector with a gold working electrode. A borate trap was used for the mobile phase followed by an amino trap for sample clean up. The analysis was performed using a Carbopac PA10 guard and analytical column. Data were acquired, based on the standard *N-*acetyl neuraminic acid peak at 5.1 min. The mobile phase was 0.1 M NaOH: 0.1 M sodium acetate (45:55) at a flow rate of 0.8 mL/min. Total run time was 30 min [31].

##### Aluminum content

Aluminum from the experimental vaccines was evaluated accordingly to European Pharmacopoeia [32].

##### Statistical analysis

The Kruskal–Wallis test was used to evaluate differences between the experimental vaccines 0 (before vaccination) and 15 days after the last dose. Differences were considered significant at p < 0.05. The ELISA and SBA data of the two strains were compared with the Mann–Whitney test. All data were analyzed using Statgraphics Plus Professional ver. 4.1 software (Statgraphics, Warrenton, VA, USA).

## RESULTS

### Physicochemical controls

The experimental formulations were sterile, more than 80% of vaccine antigens adsorbed in Al (OH)_3_, had appropriate pH, sialic acid, aluminum, and antigen concentrations in accordance with the established quantitative limits. The OMVs in residual LOS varied from 5 to 10% of the OMV protein (Table 1).

**Table 1 –.**
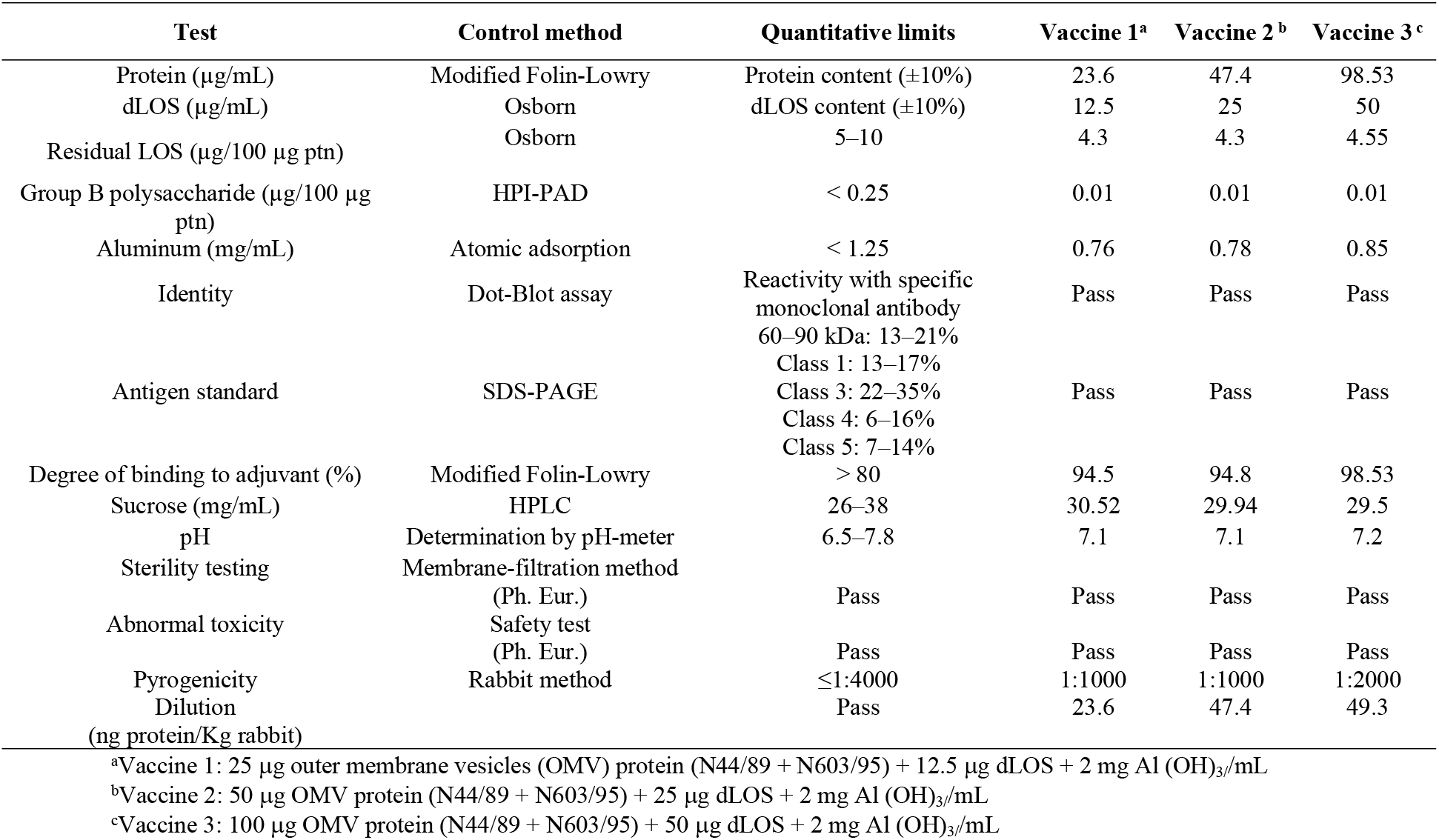
Physicochemical and biological controls for the tested vaccines.

The base peak was identified as a multiple-charge ion [M+2H-H_2_O]^+2^ at m/z 1387.9, after charge deconvolution, showed molecular weight compatible with a dihexosamine backbone, carrying 1,4’ diphosphate groups (PP) and *N*,*N*-diacylated by 3-hydroxy-tetradecanoyl fatty chains. No variation of mass was observed for OS core moiety, (Figure 1).

**Figure 1:**
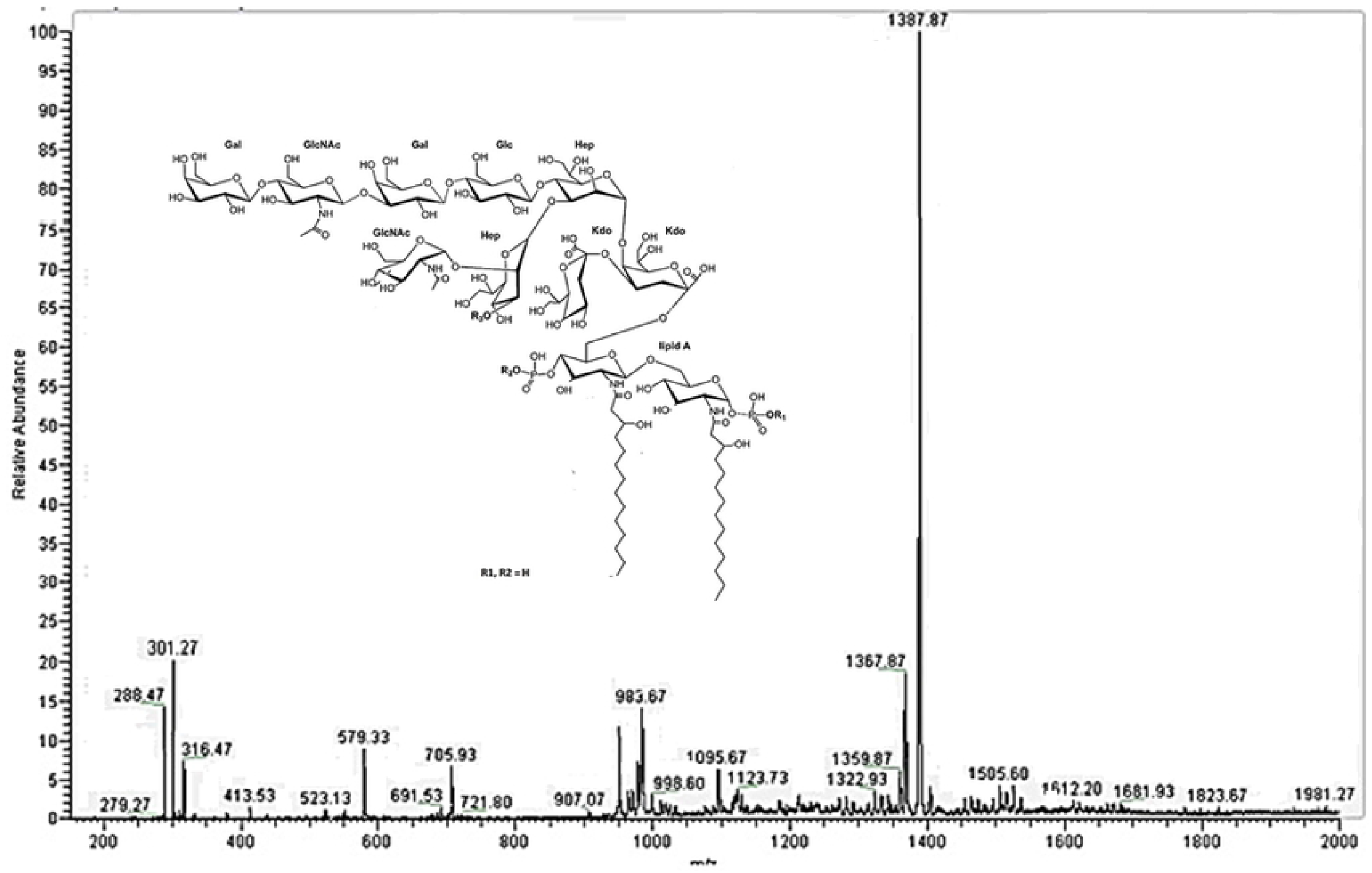
Electrospray ionization/mass spectroscopy (ESI/MS) of O-deacylated lipooligosaccharide (LOS) from N44/89 *Neisseria meningitidis* B, strain.

A second minority ion was also analyzed. A sodiated-protonated ion [M+H+Na] ^+2^ at m/z 1367.87. It had mass compatible with a diacylated lipid A and OS core discribled previously (m/z 1387.9), but differs in their phospho groups, showing only a monophosphorylated (P) form. Native LOS was also analyzed showing molecular weight compatible with and hexacylated structure carrying two units of phosphoethanolamine (data not shown).

### Morphology

The morphology of the final N44/89 preparation obtained by electron microscopy is presented in Figure 2. The OMV range in size and morphology after sterile filtration were around 80–100 nm.

**Figure 2:**
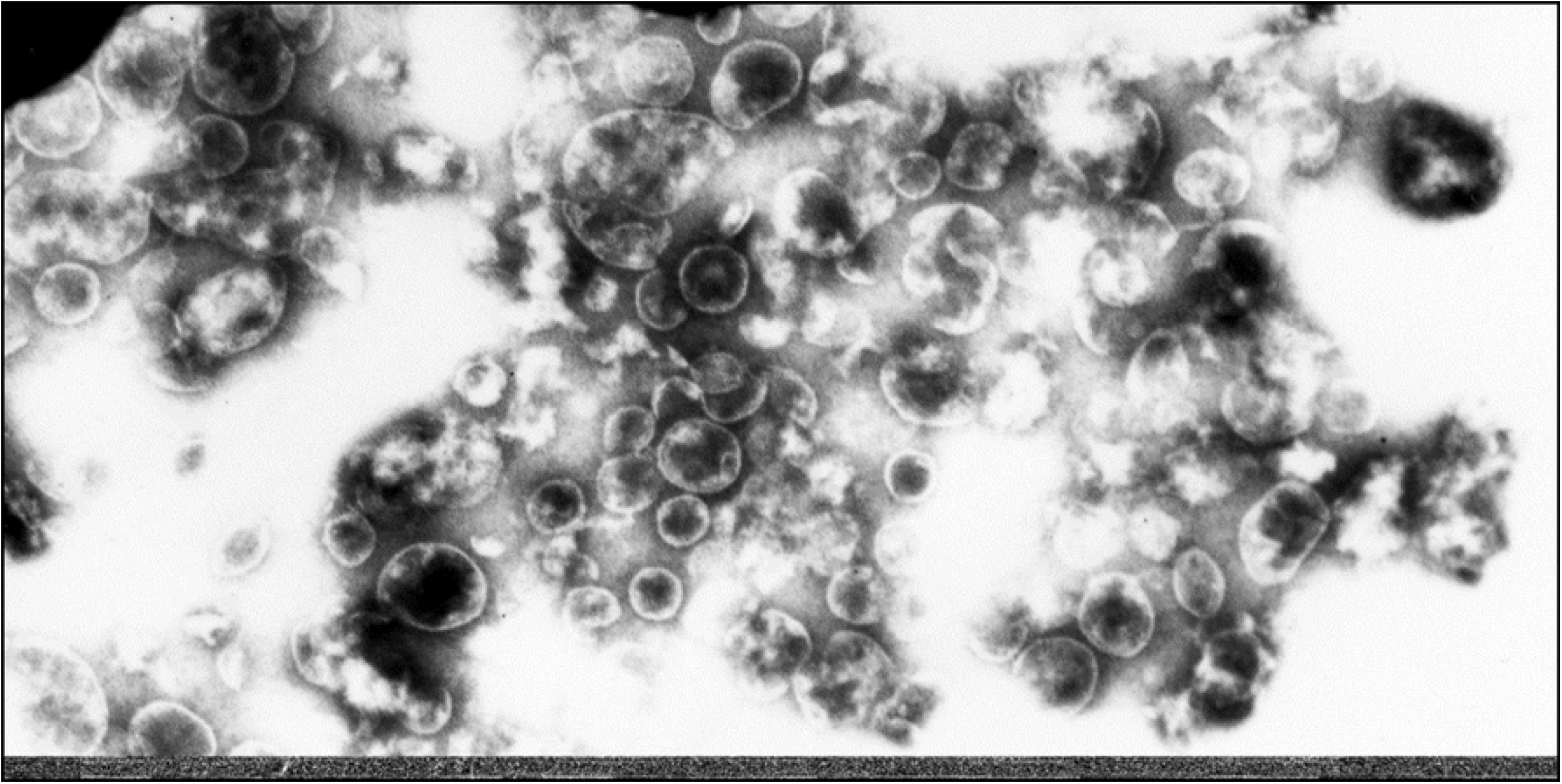
Electron microscopy of outer membrane vesicles (OMV) from the N44/89 strain (1 mg/mL) in distilled water negatively stained with 0.5% (w/v) phosphotungstic acid, pH 7.0 The preparations were examined under an electron microscope (JEM 1010; Jeol, Tokyo, Japan).

### Biological Controls

#### Vaccines immune response in mice

The three experimental vaccines induced significant increases in total IgG (p < 0.001) in response to the OMVs (Table 2) of both Brazilian prevalent strains with no difference between them. Seroconversion to LOS (300 and 1,000-fold) was detected 15 days after the third dose for vaccines 2 and 3, respectively.

**Table 2:** Total IgG geometric mean concentrations (GMCs) against vaccine components from the N44/89 and N603/95 strains before and after immunizing mice with three doses of a vaccine as determined by enzyme-linked immunosorbent assay.

| Mice total antibody by ELISA against OMVs from strains N44/89 and N603/95 and LOS from N44/89 strain |  |  |  |  |  |  |  |  |  |
| --- | --- | --- | --- | --- | --- | --- | --- | --- | --- |
| Study time | N44/89 (B:4,7:P1.19,15, L1,3,7,8)<sup>d</sup> |  |  | N603/95(B:4:P1,7,1, L,3,7)<sup>e</sup> |  |  | LOS (B:4,7:P1.19,15, L1,3,7,8)<sup>f</sup> |  |  |
|  | 25 µg /mL<sup>a</sup> | 50 µg /mL<sup>b</sup> | 100 µg /mL<sup>c</sup> | 25 µg /mL<sup>a</sup> | 50 µg /mL<sup>b</sup> | 100 µg /mL<sup>c</sup> | 25 µg /mL<sup>a</sup> | 50 µg /mL<sup>b</sup> | 100 µg /mL<sup>c</sup> |
| Day 0 | <DL | <DL | <DL | <DL | <DL | <DL | <DL | <DL | <DL |
| Before 2<sup>nd</sup><br>dose (Day 15) | 263 ± 32 | 868 ± 40 | 185 ± 40 | 113 ± 8.28 | 446 ± 44 | 448 ± 12 | <DL | <DL | <DL |
| Before 3<sup>rd</sup><br>dose (Day 30) | 3 508 ± 339 | 7027 ± 182 | 11 596 ± 649 | 4128 ± 478 | 7008 ± 215 | 11 061 ± 253 | <DL | <DL | <DL |
| After 3<sup>rd</sup> dose<br>(Day 45) | 14 946 ± 1<br>138 | 8415 ± 497 | 16 262 ± 191 | 14 368 ± 115 | 11 781 ± 203 | 10 279 ± 243 | <DL | 334.6 ± 16.5 | 1002 ± 142.3 |

<sup>a</sup>Vaccine 1: 25 µg OMV protein (N44/89 + N603/95) + 12.5 µg detoxified endotoxin (dLOS) + 2 mg Al(OH)<sub>3</sub>/mL
<sup>b</sup>Vaccine 2: 50 µg OMV protein (N44/89 + N603/95) + 25 µg dLOS + 2 mg Al (OH)<sub>3</sub>/mL
<sup>c</sup>Vaccine 3: 100 µg OMV protein (N44/89 + N603/95) + 50 µg dLOS + 2 mg Al (OH)<sub>3</sub>/mL
<sup>d</sup>OMV from N44/89 as coating antigen; <sup>e</sup>OMV from N603/95; LOS<sup>f</sup> as coating antigen.
<sup>g</sup>N<sup>o</sup> of subjects: three pools of five mice each in the indicated group: GMC ± standard error; NE – non-evaluated; <DL below detection limit.

Vaccine 1 did not induce a LOS response in mice at the same evaluation time, suggesting that the immune antigen response was concentration dependent. Considering a fourfold increase in bactericidal titres, after 3 doses of vaccines as a immunogenicity parameter, the experimental vaccines were very immunogenic in mice. The response for N44/89 strain was stronger than for N603/95 even in the vaccine with the lowest concentration of vaccine components. Based on the bactericidal activity of pre immune sera, the response to N44/89 was 4 and 8 times higher than N603/95 in the intermediate and maximum antigen vaccines concentration, respectively. In addition, the bactericidal titers were significantly higher for N44/89 than what was observed to N603/95 after the third dose (Table 3).

**Table 3:** Serum bactericidal assay (SBA) antibody titers (reciprocal serum dilution ≥ 50% killing) against group B vaccine strains before and after immunizing mice with a three-dose vaccination schedule using phase II trial meningococcal experimental vaccines.

| Bactericidal titers indicated as reciprocal sera dilution $\geq 50\%$ killing<sup>a</sup> | | | | | | |
| --- | --- | --- | --- | --- | --- | --- |
| Study time point | N44/89 (B:4,7:P1.19,15:P5.5,7: L1,3,7,8)<sup>e</sup> |  |  | N603/95(B:4:P1,7,1: P5.5,7: L,3,7)<sup>f</sup> |  |  |
| Experimental<br>Vaccines | 25 µg /mL<sup>b</sup> | 50 µg /mL<sup>c</sup> | 100 µg /mL<sup>d</sup> | 25 µg /mL<sup>b</sup> | 50 µg /mL<sup>c</sup> | 100 µg /mL<sup>d</sup> |
| Pre-immune sera<br>(Day 0) | 2 | 2 | 2 | 2 | 2 | 2 |
| After 3<sup>rd</sup> dose<br>(Day 45) | 256<br>(128–256) | 256<br>(128–256) | 512<br>(256–512) | 2<br>(2–2) | 64<br>(32–64) | 64<br>(32–64) |

<sup>a</sup>Sera from three-five pools of five mouse (GMT)
<sup>b</sup>Vaccine 1: 25 µg OMV protein (N44/89 + N603/95) + 12.5 µg detoxified endotoxin (dLOS) + 2 mg Al (OH)<sub>3</sub>/mL
<sup>c</sup>Vaccine 2: 50 µg OMV protein (N44/89 + N603/95) + 25 µg dLOS + 2 mg Al (OH)<sub>3</sub>/mL
<sup>d</sup>Vaccine 3: 100 µg OMV protein (N44/89 + N603/95) + 50 µg dLOS + 2 mg Al (OH)<sub>3</sub>/mL
<sup>e</sup>N44/89 as SBA target strain.
<sup>f</sup>N603/95 as SBA target strain

#### toxicity and pyrogenicity

Both mice and guinea pigs survived to 7 days of time observation without any sign of toxicity, such weight loss, due to experimental preparations and reference vaccines administration. According to the WHO requirements for polysaccharide vaccines against *N. meningitidis*, the final product should not induce fever in rabbits at a 1:4,000 dilution of the human dose. Vaccines with 25 and 50 μg protein/mL were approved in rabbits when the human dose was diluted to 1:1,000. A vaccine with 100 μg protein/mL was approved in rabbits diluted 1:2,000 of the human dose. Cuban and DTP vaccines were approved in 1:4000 of human dose. In terms of minimum pyrogenic dose based on protein amount, the experimental preparations 1, 2 and 3, were respectively 23,6, 47,4 and 49,3 ng of protein per kg of rabbit. The reference vaccines were 12,5 ng of protein per kg of rabbit for VAMENGOC-BC^®^ and the DTP vaccines. The experimental vaccines were less pyrogenic than the references. Raw and dLOS, were tested in the rabbit pyrogen test at the highest quantity of the third (50 μg/mL) experimental vaccine. After a 30 min NaOH treatment, a sharp fall in pyrogenicity of the raw LOS was detected. The dLOS samples from the 120 and 150 min treatments were approved by the rabbit pyrogen test at dilutions of 1:200 and 1:100, respectively (Table 4).

**Table 4:**
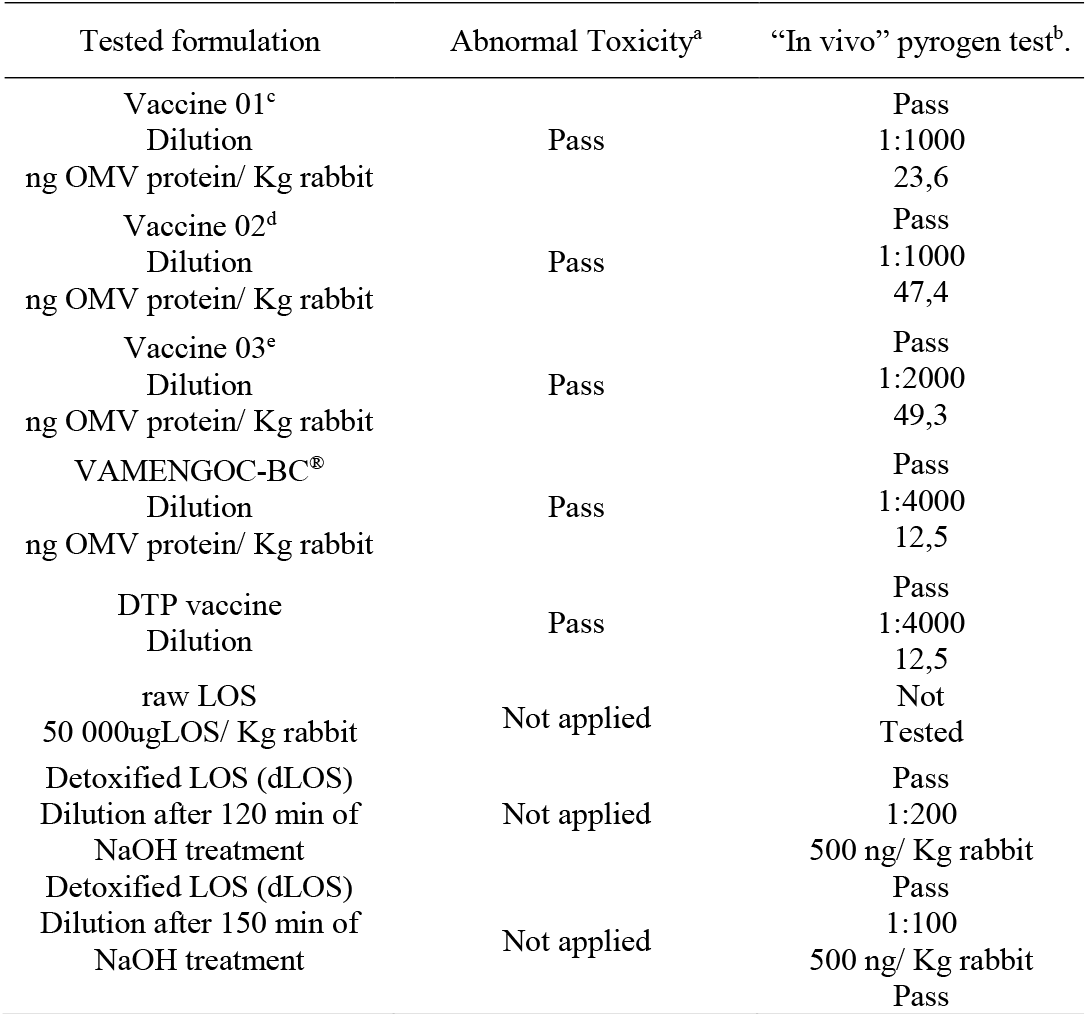
Vaccines toxicity and “in vivo” pyrogen test.

#### protein and LOS eletrophoretic patterns

The iron starvation strategy for vaccine strain growth induced IRP expression in OMV outer membrane from both vaccine strains in the molecular weight range between 60 and 90kDa. Por A and other class proteins were well expressed in OMV from N44/89 and N603/95 (Figure 3).

**Figure 3:**
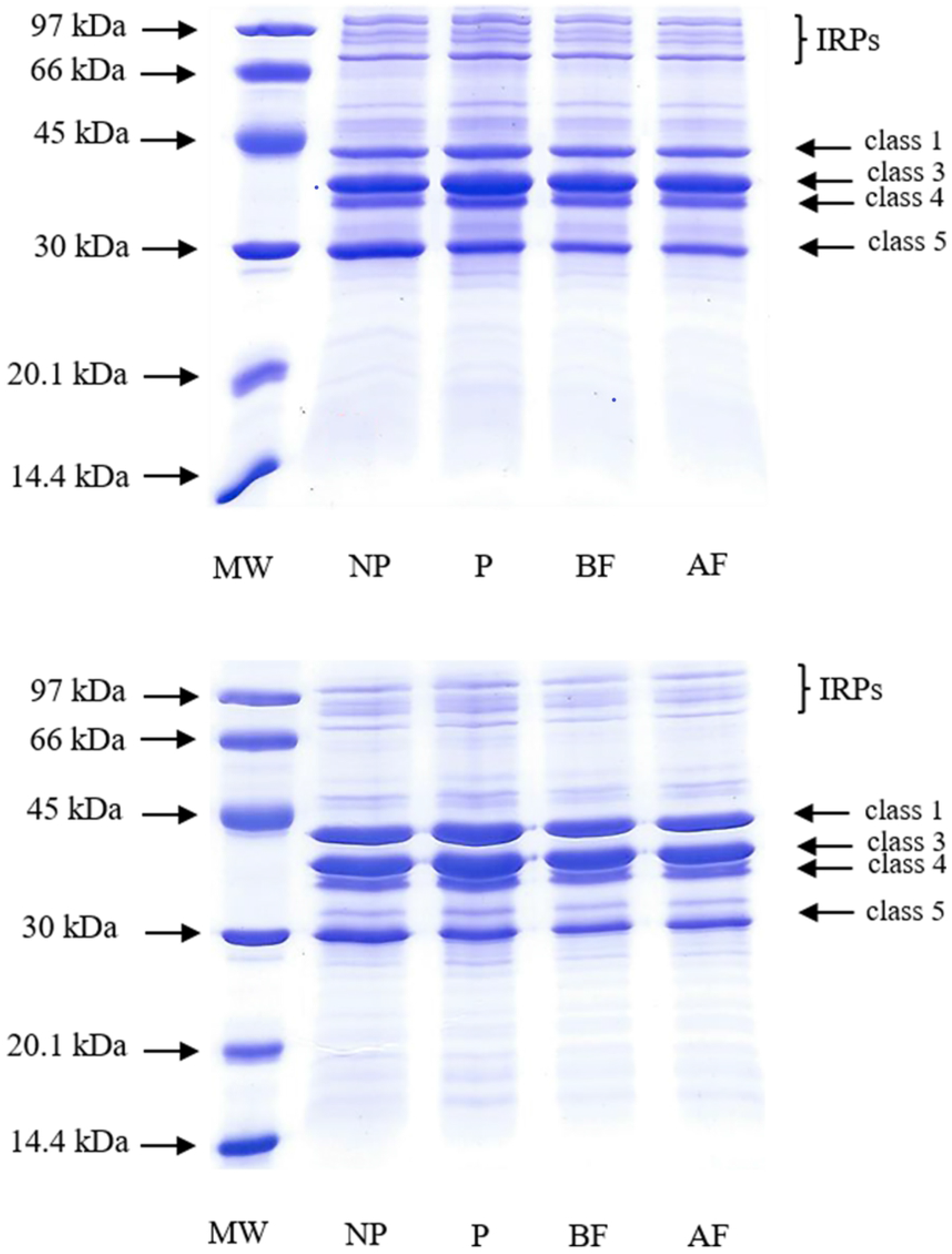
Sodium dodecyl sulfate-polyacrylamide gel electrophoresis (10 μg/outer membrane vesicle [OMV] protein) protein profile of vaccine strains OMV (A) N603/95 and (B) N44/89. Molecular weight (MW); (NP) OMV before sodium deoxy cholate (DOC) treatment; (P) OMV after DOC treatment; (BF) vaccine bulk before sterile filtration; (AF) vaccine bulk after sterile filtration.

Table 5 shows the percentages of expression of the main OMV proteins of the vaccine strains used as reproducibility parameter of the production process of this vaccine component.

**Table 5:** Class protein proportion (%) of vaccine bulk after sterile filtration by densitometry

| Protein (%) | N44/89 | N603/95 |
| --- | --- | --- |
| Por A | 23.54 | 8.10 |
| Por B | 28.84 | 15.26 |
| Class 4 | 7.63 | 1.94 |
| Class 5 | 15.95 | 9.67 |

The structural changes in Lipid A from raw LOS after the NaOH treatment are shown in the SDS profiles. Both molecules had two bands but the molecular weights of dLOS were lower than those shown in LOS profile before alkali detoxification (Figure 4).

**Figure 4:**
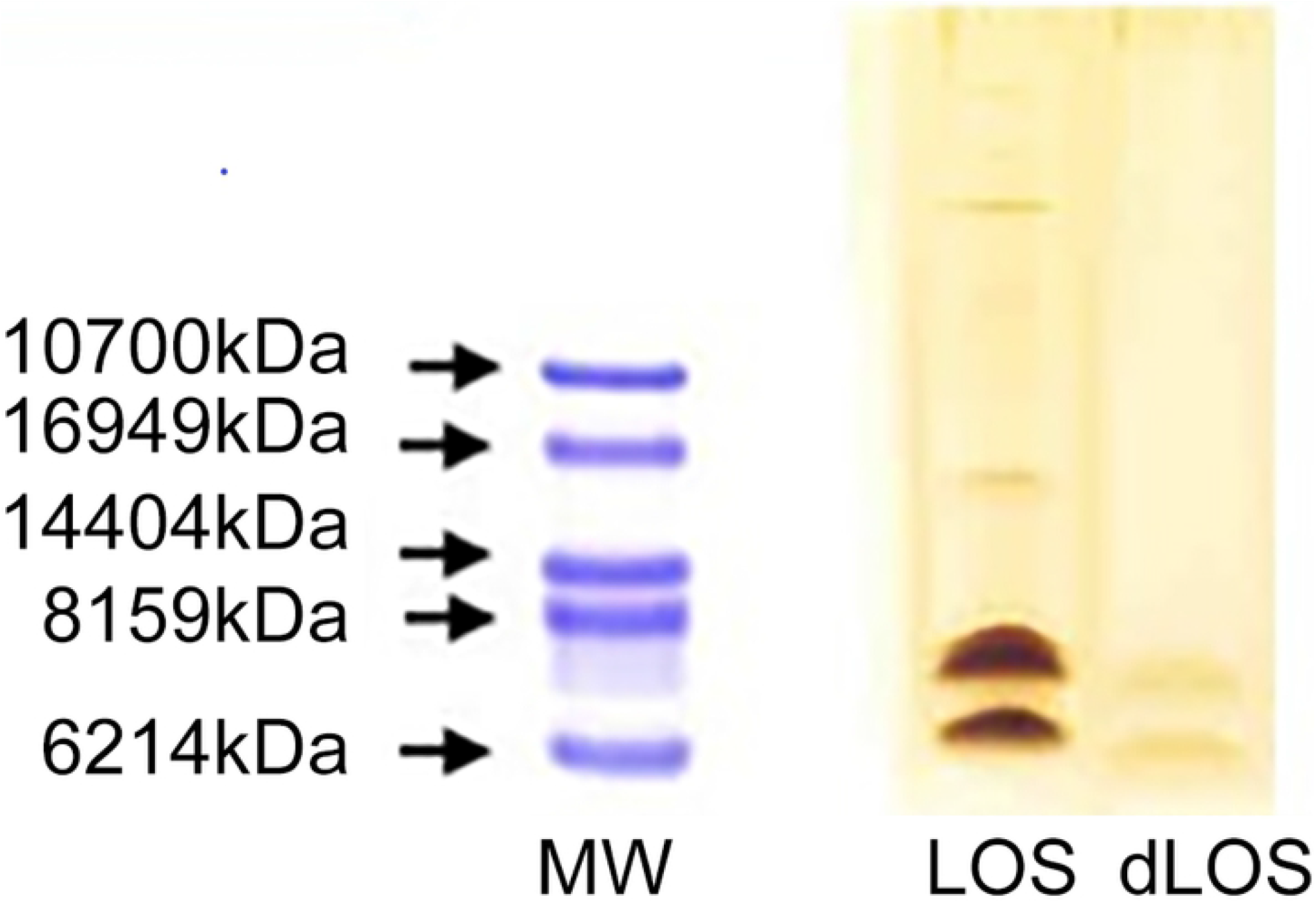
Sodium dodecyl sulfate-polyacrylamide gel electrophoresis lipooligosaccharide (LOS) profile from N44/89 biomass. MW, protein molecular weight standards of 16.949–2.512 kDa; LOS, native endotoxin; dLOS, detoxified endotoxin.

## DISCUSSION

The poor immunogenicity of serogroup B polysaccharide has focused on the study of subcapsular antigens as an alternative approach to have an effective vaccine against meningococcal disease caused by this serogroup [10,11, 33].

In countries in which meningococcal disease is caused by a specific strain of meningococcus B, OMV vaccines have been shown to be effective. Their serotype specificity restricts the induction of cross-immunity among the wide variety of pathogenic strains of this serogroup indicating the preferred use of these products for outbreaks control [34].The OMV vaccines induce protective antibodies mainly against Por A, considered their major immunodominant antigen, which is the main target for SBA activity. The vaccine high efficacy is dependent of the similarity between Por A from vaccine and circulating strains. However, minor antigens as other outer membrane proteins and LOS, acting synergistically with Por A can contribute with vaccine efficacy [35,36]. The use of native OMV conformation as vaccine component, requires a detergent treatment to decrease LOS concentration aiming its decrease toxicity. LOS is an additional subcapsular antigen in OMV and has been studied to broaden coverage and cross-reactivity of serogroup B meningococcal vaccines (zolliner).Despite successfully decreasing toxicity, the outer membrane components that contribute to induce the immune response can also be affected. It was very important the establishment of all physico-chemical controls of vaccine components according to the literature, that will presented to Brazilian regulatory agency to go forwards in clinical trials (Table 1),.

The Brazilian vaccine against *N.meningitidis* serogroup B has in its composition, outer membrane vesicles (OMV) from Brazilian prevalent strains. They were obtained from bacteria growth with free iron starvation using ethylene diamine-di-(O-hydroxy-phenylacetic acid) (EDDHA) as iron chelator. This approach aimed to simulate the condition of availability of free iron in the blood of human host. The EDDHA chelating and release iron constants are similar to transferrin constants, which is the protein carrier of this element, in the blood. The presence of iron attached to protein carriers or EDDHA in bioreactor, induces the expression of iron regulated proteins (IRPs) on *N.meningitidis* OMV membrane that may be important in vaccine-induced immune response and increase of bacteria pathogenicity. Outer membrane from *B.pertussis* obtained from growth under iron starvation conditions, showed higher expression of proteins regulated by this ion as well as decrease in LPS phosphorylation degree compared to standard molecule [37, 38, 39,40].

The mass spectrometry data showed a severe reduction of mass comparing native and detoxified LOS. The observed mass variation was attributed to the hydrolysis of fatty acyl esters and partial hydrolysis of phospho groups, which generated a less toxic molecule (Figure 1).

Detergent-treated OMVs mainly induce immune response to parental strains homologous to Por A, which provides limited cross-protection of these vaccines to different serosubtypes and serogroups. The expectation of an increase in vaccine cross-reactivity using LOS as additional component was based on previous studies. The L3, 7, and 8 Brazilian prevalent strain immunotypes are expressed on a high percentage of serogroup B strains associated with epidemics [41]. The expectation of an increase in vaccine cross-reactivity using LOS as additional component must be evaluated in human clinical trials of the developed vaccine [42]. A significant increase in total IgG to OMV was observed, in mice, for both vaccine strains after immunization with the three experimental preparations, compared to pre immune values. It can be used as a primary vaccine potency control parameter. There was no difference between three vaccine concentrations. Total IgG produced against LOS increased 15 days after the third dose only for the most concentrated vaccines. Moreover, the increase was concentration-dependent, as the highest LOS concentration induced the highest total IgG concentration. Total IgG produced against LOS did not indicate a different role for residual LOS on OMVs and added dLOS in the vaccine immune response (Table 2). The role of each of these molecules in inducing a protective response against meningococcus should be evaluated in human clinical studies.

However the use of LOS as vaccine component, demands de decrease of its toxicity not to increase OMV intrinsically pirogenicity. A NaOH treatment is required to remove the ester-linked fatty acid tails from lipid A, which are responsible for the toxicity. dLOS as a potential target of serogroup B meningococcal vaccines is based on its contribution in bactericidal and opsonic activities of induced antibodies in meningococcal disease and after immunization with OMV vaccines [36, 37,38]. The samples of LOS and dLOS were analyzed by SDS-PAGE, presenting difference in electrophoretic profile. The difference observed was attributed to Lipid A reduction degree of acylation, after NaOH treatment, which was confirmed by ESI-MS analysis (Figure 2).

The SBA titers to N44/89 induced in mice were protective after the first dose and remained protective for 15 days after the third dose of all preparations with no increase after the third dose. On the other hand, the SBA titers to the N603/95 strain were protective only after the second dose, suggesting that the third dose was important. In addition, the N603/95 titers were significantly lower than for N44/89 using all vaccines. The values remained at lower levels for 15 days after the third dose (Table 3).

Human and murine-specific protecting antibodies are mainly directed against two long, hypervariable, extracellular regions in Por A loops 1 and 4. However, other minor proteins and LOS are antigens in the total immune response induced by OMV vaccines. Although Por A is the dominant antigen, other outer membrane components may act synergistically during the induced immune response against meningococcal B. Antibody affinity maturation was induced in response to the dominant antigen but was seriously delayed in response to a subdominant antigen, resulting in less functional serum IgG. The expression of minor proteins and their localization in the outer membrane are mandatory to activate the complement cascade via the classical pathway, which is the first step to eliminate the bacteria [35]. The Por A from N603/95 (P1.7.1) may have been a subdominant antigen compared to that of P1.19.15 from N44/89 during the immune response to the experimental vaccines, which may contributed to SBA lower titres to N603/95 strain. The weaker immune response observed to the N603/95 strain may have been caused by weaker or less accessibility of B cell epitopes due to physical and chemical properties, such as hydrophobicity, length, or the combination of OMVs from the vaccine strains [21].

The Brazilian vaccine development approach, uses multiple antigens from B prevalent strains with the aim of broading vaccine coverage (Figure 3 and 4). The combination of prevalent vaccine strains with different PorA (P1.19.15 and P1.7.1 from N44/89 and N603/95 respectively), and LOS immunotypes 3, 7, and 8, shared among different B strain serosubtypes followed the tailor-made vaccine concept based on Brazilian meningococcal disease epidemiology [41].

The rabbit pyrogen test has been used historically to measure pyrogenicity potential of human and animal vaccines. However, whole cell and OMV based vaccines, which intrinsically contain relatively high concentration of pyrogenic material, has been a source of great concern. Several studies have been reported to define the best biological tests that can be used as safety parameters for clinical trial [43].

The pyrogenicity evaluation of OMV based vaccines has been shown that the way of production of these structures as well as the characteristics of the rabbit’s strains can affect the test response what has hampered the establishment of the acceptable safety range for new vaccines, in the pyrogen test “in vivo”. Although the results showed here are based on protein concentration, the minimum pyrogenic dose established in pirogenio test “in vivo” considers all pyrogens. In this way, to experimental vaccines are being considered all vaccine components contributing to the “in vivo” pyrogen test results [9,10].

Experimental vaccines showed minimum pyrogenic dose between 23 and 50 ng of protein per rabbit body weight. The differences observed for the vaccine with 25ug and 50ug of OMV protein /mL can be attributed to inherent “in vivo” pyrogenic test variation. Although the vaccine with the highest concentration (100ug/mL protein), also presented 50ng ng of protein per rabbit body weight as minimum pyrogenic dose, it was achieved with double dilution compared to vaccines with lower concentration of OMV protein. The results suggested that OMV concentrations higher than 50ng of protein per kg of rabbit body weight, the pyrogenicity activity seems to concentration dependent (Table 4).

Most of OMV based vaccines clinical studies, used 50 or 100 μg of OMV protein/mL but none of them, combining dLOS and OMVs from two strains, as the Brazilian B vaccine [33,34]. In the Brazilian developed vaccines, residual LOS from OMVs, plus dLOS from the N44/89 strain composed the total amount of LOS. To assess the contribution of dLOS in experimental preparations, it was also evaluated in the pyrogenic test “in vivo”. The test performed with raw and detoxified LOS showed a decrease of 1×10^6^ in LOS pyrogenicity activity which may attributed to de Liped A desacetillation [9,10].

In the development of the Brazilian vaccine against meningococcus B, as for other producers, we used as a reference the WHO requeriments for meningococcal polysaccharide vaccines. The vaccine should be approved in rabbit pirogen test, in the dilution of 1:4000 of human-dose or 25 ng polysaccharide/rabbit kg body weight as the minimum pyrogenic dose of *N.meningitidis* polysaccharide. According to what was established by the WHO, all tested vaccines would be safe for clinical trials in humans as they have passed the pyrogenicity test in rabbits at dilutions less than 1: 4000 of the probable human dose.The OMV-based vaccine, VAMENGOC-BC^®^, and the DTP vaccine officially included in Brazilian Immunization Program calendar were used as references in the “in vivo” pyrogenic test to compare the experimental vaccine pyrogenic potential. Both vaccines, would be approved by WHO for clinical trials. However, their potential pyrogenicity were higher than what was observed for the three experimental vaccines in rabbits [44].

The production approach applied to obtain OMVs from Brazilian prevalent strains was not much different from that described for OMV vaccines. The experimental preparations were approved by rabbit pyrogen test results of < 20,000 EU/mL vaccine. Residual LOS toxicity in the OMVs was 5–10 μg/100 μg total protein, which is in accordance with previous studies [45,46, 47].

These study results indicate that the use of the Brazilian meningococcal B tested vaccines are in accordance with the literature for similar products and clinical trials studies can be considered.

## CONFLIT OF INTEREST

All authors have no conflict of interest to declare.

## ACKNOWLEDGMENTS

We would like to thank Dr. Carl Edward Frasch and Dr Akira Homma for their support and guidance. This manuscript would not be possible without your contributions.

## AUTHOR CONTRIBUTIONS

All authors contributed equally in the elaboration of this article.

## Supporting information

S1 - Figure 1: Electrospray ionization/mass spectroscopy (ESI/MS) of O-deacylated lipooligosaccharide (LOS) from N44/89 *Neisseria meningitidis* B, strain.

**(File: Figure 1 PLos One Ellen Jessouroun.tif)**

S2 - Figure 2: Electron microscopy of outer membrane vesicles (OMV) from the N44/89 strain (1 mg/mL) in distilled water negatively stained with 0.5% (w/v) phosphotungstic acid, pH 7.0 The preparations were examined under an electron microscope (JEM 1010; Jeol, Tokyo, Japan).

**(File: Figure 2 PLOs One Ellen Jessouroun_300dpi.tif)**

S3 - Figure 3: Sodium dodecyl sulfate-polyacrylamide gel electrophoresis (10 μg/outer membrane vesicle [OMV] protein) protein profile of vaccine strains OMV (A) N603/95 and (B) N44/89. Molecular weight (MW); (NP) OMV before sodium deoxy cholate (DOC) treatment; (P) OMV after DOC treatment; (BF) vaccine bulk before sterile filtration; (AF) vaccine bulk after sterile filtration.

**(File: Figure 3 PLOs One Ellen Jessouroun .tiff)**

Figure 4: Sodium dodecyl sulfate-polyacrylamide gel electrophoresis lipooligosaccharide (LOS) profile from N44/89 biomass. MW, protein molecular weight standards of 16.949–2.512 kDa; LOS, native endotoxin; dLOS, detoxified endotoxin.

**(File: Figure 4 PLOs One Ellen Jessouroun.tiff)**

